# Novel Computed Tomography-based tools reliably quantify plant reproductive investment

**DOI:** 10.1101/182386

**Authors:** Yannick M. Staedler, Thomas Kreisberger, Sara Manafzadeh, Marion Chartier, Stephan Handschuh, Susanne Pamperl, Susanne Sontag, Ovidiu Paun, Jüerg Schöenenberger

## Abstract

The flower is a bisexual reproductive unit where both genders compete for resources. Counting pollen and ovules in flowers is essential to understand how much is invested in each gender. Classical methods to count very numerous pollen grains and ovules are inefficient when pollen grains are tightly aggregated, and when fertilization rates of ovules are unknown. We thus established novel, Computed-Tomography-based counting techniques. In order to display the potential of our methods in very difficult cases, we counted pollen and ovules across inflorescences of deceptive and rewarding species of European orchids, which possess both very large numbers of pollen grains (tightly aggregated) and ovules. Pollen counts did not significantly vary across inflorescences and pollination strategies, whereas deceptive flowers had significantly more ovules than rewarding flowers. The within inflorescence variance of pollen to ovule ratios in rewarding flowers was four times higher than in deceptive flowers, possibly demonstrating differences in the constraints acting on both pollination strategies. We demonstrate the inaccuracies and limitations of previously established methods, and the broad applicability of our new techniques: they allow measurement of reproductive investment without restriction on object number or aggregation, and without specimen destruction.

## Introduction

It should be evident to most human beings that the mere existence of genders is a harbinger of conflicts for resources. Gender conflicts are nowhere more acute than in hermaphroditic organisms where both genders have to draw from the resource pool of the same organism to maximize fitness (Charnov, 1979; Lloyd, 1979). In the overwhelmingly hermaphroditic flowering plants, counting the pollen grains and ovules of flowers allows us to understand how much a plant invests in the male vs. female part of its fitness.

Pollen counting methods fall into three groups (Costa and Yang, 2009): counting with the naked eye, particle counters, and image processing algorithms. Counting visually usually involves spreading samples on specialized slides with a grid, and counting a sub-sample (e.g. (Jorgensen, 1967) (Kearns and Inouye, 1993)), which is then extrapolated (Kannely, 2005). Pollen grains tend to settle disproportionately on grids, especially if the grains are large or still aggregated, which may then produce incorrect estimates (Kannely, 2005). Electronic or laser-based counters physically detect pollen grains in order to count them. A particle counter counts every particle; unfortunately, this may include debris and aggregated pollen (Kearns and Inouye, 1993). Image processing automates pollen counting from pollen grain images. This requires software to scan images for objects and then count each object as a unit (e.g. (Bechar *et al.*, 1997), (Aronne *et al.*, 2001)). All three approaches require sample destruction, including proper de-aggregation of the grains, which can be hard to achieve.

Ovules are counted either before fertilization (as ovules) or after fertilization and maturation (as seeds). At the ovule stage, counting is manual after dissection, either directly or on photographs (Nazarov, 1989), either on the whole ovary or an a stretch of the latter (ovules per mm are then extrapolated to the length of the whole gynoecium). At the seed stage, methods fall into two groups: (1) extrapolations based on counting an aliquot of known weight (Salisbury, 1942), or on a portion of a line of dry seeds (Darwin, 1862), or on the surface of a liquid (Burgeff, 1936), or within a suspension of seeds (Proctor and Harder, 1994; Sonkoly *et al.*, 2016). (2) use of particle counting devices (DuBois, 2000). All these approaches require sample destruction and rely on the assumption that all ovules have been fertilized, which is hard to test.

In summary, these traditional methods for counting pollen and ovules show limitations with very large numbers of pollen grains (that are tightly aggregated) and ovules (especially when fertilization rates are unknown). Given these limitations, novel and more reliable methods to count pollen and ovules are needed. Contrasting agents for X-Ray Computed Tomography (CT), especially phosphotungstic acid (PTA), semi-selectively accumulate in both pollen and ovules, possibly due to the higher protein content of their cells/tissues relative to their surroundings (Bellaire *et al.*, 2012; Hayat, 2000; Staedler *et al.*, 2013). By selecting only the brightest voxels (3D pixels) of the 3D model, i.e., the areas that absorb the more X-Rays (X-Ray data is traditionally displayed in negative), it is possible to segregate pollen and ovules from their surroundings tissues. This process is called greyscale thresholding. Obtaining the volume of such a selection by counting voxels is straightforward. Provided that the average volume of a grain or ovule is available, CT can thus be used to count ovules and pollen in case where classical methods are at their limits.

These limits are nowhere more evident than in the study of orchids, which both possess enormous numbers of ovules and –tightly aggregated-pollen grains (Darwin, 1862). Orchids are remarkable plants in many ways: not only are they the largest family of flowering plants (with up to 30 000 species), but also almost a third of them offer no reward to their pollinators (Ackerman, 1986; Porsch, 1909; Van der Pijl and Dodson, 1966). The presence of a reward, or lack thereof, influences dramatically pollinator behaviour: in rewarding taxa, the pollinators tend to visit several or all open flowers of an inflorescence during a visit, and visit the same inflorescence repeatedly in order to harvest its rewards ((Van der Cingel, 1995); Fig. 1A, B). In taxa with deceptive flowers, however, pollinators tend to learn to avoid deception and to visit only the first open flowers they encounter while they are still naive ((Jersáková and Kindlmann, 1998; Van der Cingel, 1995); Fig. 1D). Consequently, in rewarding plants, the fruit set is usually higher and fruits are spread across the inflorescence ((Neiland and Wilcock, 1995); Fig. 1C), whereas in deceptive plants, only the first flowers to open tend to bear fruits ((Jersáková and Kindlmann, 1998; Nilsson, 1980; Vogel, 1993); Fig. 1E). In orchids in general, the ratio of pollen to ovules (P:O) increases from the bottom to the top of inflorescences (Kopylov-Gus’ kov Yu *et al.*, 2006; Nazarov and Gerlach, 1997; Salisbury, 1942). Due to decreased pollinator visits to top flowers, we hypothesise that the decrease in ovule number (increase in P:O) should be stronger across inflorescences of deceptive flowers than across inflorescences of rewarding flowers (Fig. 1F-I). The Orchidinae, to which most European orchids belong, are a good system to test this hypothesis because their pollination biology and phylogenetic relationships are very well understood (Claessens and Kleynen, 2011; Inda *et al.*, 2012; Van der Cingel, 1995). Known phylogenetic relationship between studied species is needed in order to control for potential phylogenetic constraints.

**Fig. 1.**
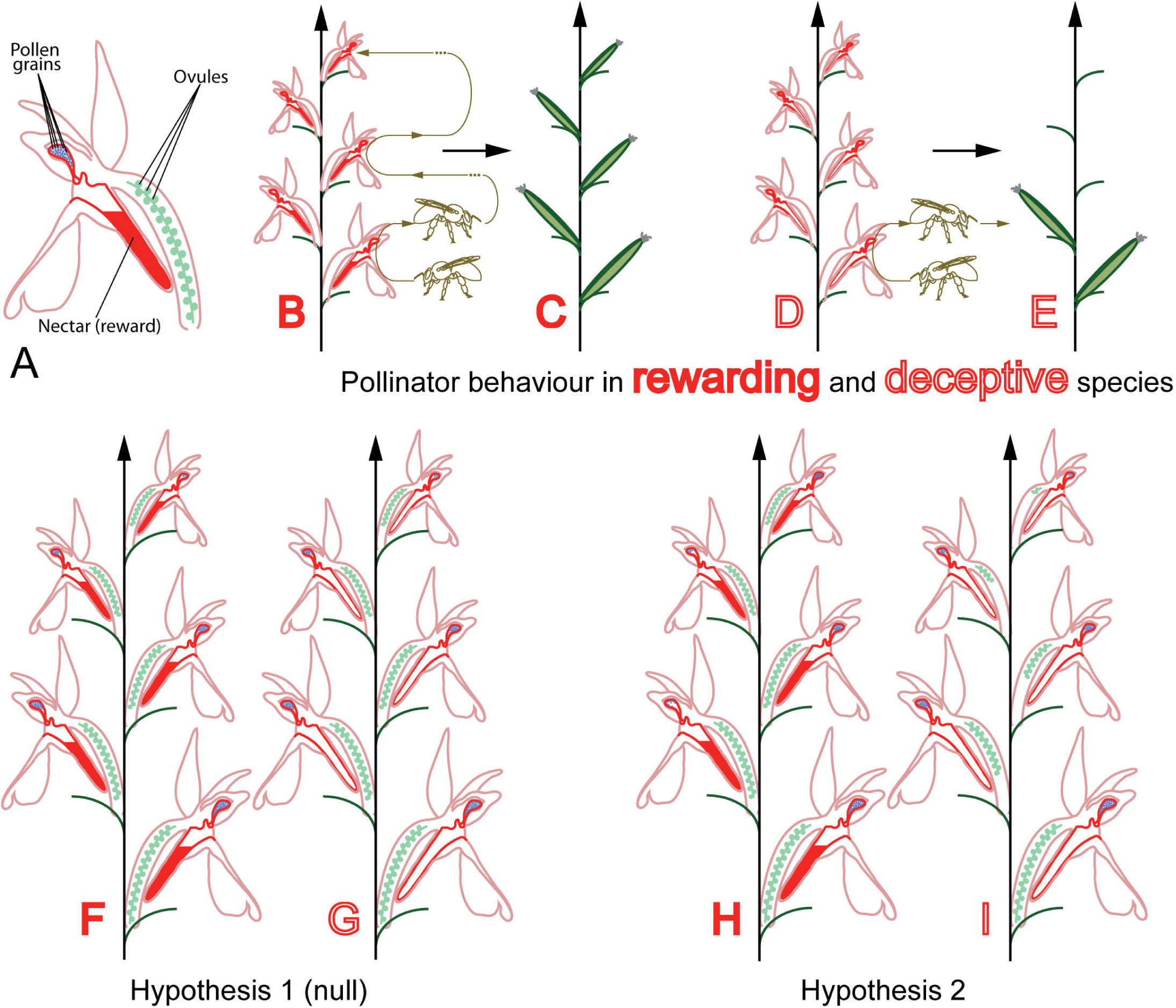
Pollinator reward or deception lead to different fruit set - hypotheses on reproductive investment. A, legend. B, schematic behaviour of pollinator on rewarding inflorescence: pollinator learns to associate the flowers with reward and visits the inflorescence repeatedly. C, fruit set in rewarding inflorescences is high equally distributed on the inflorescence. D, schematic behaviour of pollinator on deceptive inflorescence: naive pollinator learns to avoid inflorescence. E, fruit set on deceptive inflorescence is concentrated on the first flowers to open, at the bottom of the inflorescence. F, G, hypothesis 1: there is no difference between gender allocation strategy in deceptive and rewarding inflorescences. H, I, hypothesis 2: there is a difference between gender allocation strategies of deceptive and rewarding inflorescences. The difference in reproductive investment between lower and higher flowers is stronger in deceptive inflorescences than in rewarding inflorescences.

This study aims at (1) establishing new methods for pollen and ovule counting that can be used even for flowers with many, densely aggregated pollen grains and ovules, and (2) demonstrating the potential of these methods by focussing on species of European orchids to uncover if the differences of pollinator behaviour in rewarding *versus* deceptive plants lead to different patterns of reproductive investment at the level of the inflorescence.

## Materials and methods

### Material

We sampled three rewarding and five deceptive species of the subtribe Orchidinae ((Inda *et al.*, 2012); see table S1). We collected three flowers for two to four inflorescences per species, for a total of 76 flowers (table S2).

### Collection method

Open flowers (including pollinia, i.e. the pollen aggregates of orchids) or buds close to anthesis were collected from the bottom, middle and top sections of inflorescences and were immediately fixed in 1% phosphotungstic acid in formalin-acetic acid-alcohol (1% PTA / FAA; Fig. 2A). The flowers were removed from the plants with razor blades and tweezers.

**Fig. 2.**
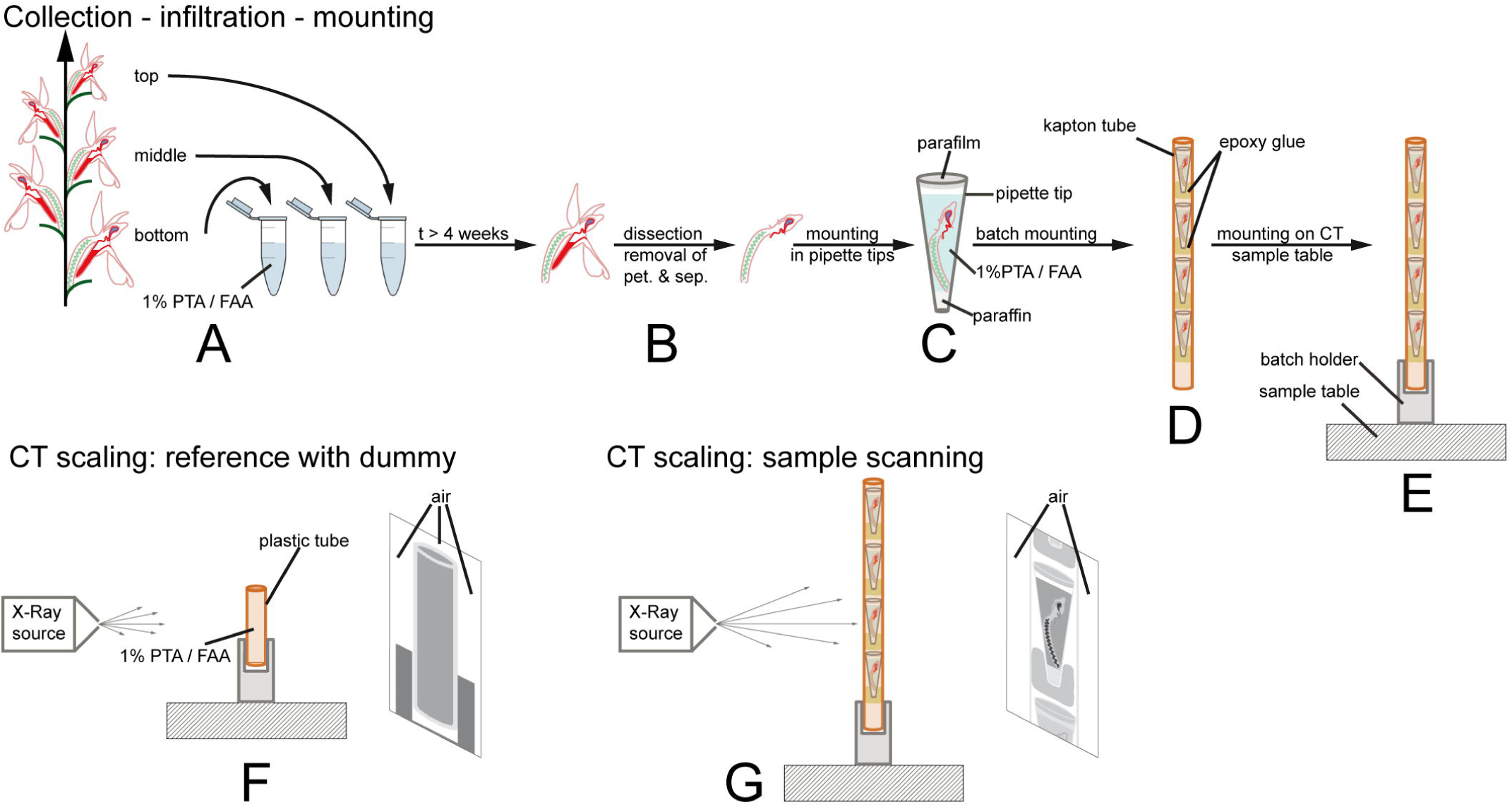
Sample processing and scanning approach. A, collection and fixation of flowers. B, removal or perianth organs of flowers. C, mounting in pipette tip. D, mounting in batches in kapton tubes. E, mounting on sample table for scanning. F, scanning of dummy for CT scaling, in order to obtain calibrated greyscale values (air has to be in the field of view on both sides and on top of the sample). G, sample scanning. Air has to be present on both sides of the sample for CT scaling to work.

### Sample preparation

The sampled flowers and buds were de-aerated for 20-30 minutes with a water-jet vacuum pump. Total infiltration time in 1% PTA / FAA was four to six weeks during which the solution was changed twice. The long infiltration time allowed saturation of the sample with contrasting agent. In order to further optimize for space constraints during mounting for CT-scanning and to prevent the formation of air bubbles, the petals and sepals were removed with tweezers and dissecting scissors (Fig. 2B). The mounting was then performed in a 200µl pipette tip (Standard UNIVERSAL (Art. No.: B002.1), Carl Roth GmbH+Co KG) as described in (Staedler *et al.*, 2013) with the following differences: (1) before mounting, the bottom end of the pipette tip was cut off. This allowed us to enter the tip with a preparation needle and to drag down the samples to optimize the used space without displacing or breaking pollinia. (2) the samples were not washed with 70% EtOH before scanning, but immediately mounted in PTA / FAA (Fig. 2C). The singly mounted flowers were then batched in longitudinally slit tubes of kapton (an X-Ray lucent material), see Fig. 2D (Ø: ca. 3mm, American Durafilm Co., Inc.). Although the tension produced by the shape of the tubes stabilized the samples, they were additionally stabilized by gluing them to the straw with epoxy glue (UHU Plus, UHU GmbH & Co. KG). The batch was then placed in an in-house batch holder and fixed with epoxy glue, itself fixed to the sample table (see Fig. 2E). Epoxy glue was then added between all the abovementioned parts to stabilize them. In this way, we were able to programme and scan batches of up to five flowers sequentially.

### Scanning

Scanning was performed with a MicroXCT-200 system (Zeiss Microscopy). Scanning conditions are summarised in table S2. 3D reconstructions were performed *via* the software XMReconstructor 8.1.6599 (Zeiss Microscopy). In order to have repeatable greyscale values from one scan to another, byte scaling and CT scaling were used. Byte scaling is a procedure in which minimum and maximum greyscale values are set for the whole reconstructed scan volume (Xradia, 2010). Byte scaling was used for the reconstruction of scans of gynoecia. CT scaling is a procedure by which greyscale values are scaled to the values of two reference materials (Candell, 2009); in our studies, air and a solution of 1% PTA in FAA were used. CT scaling requires scanning of a dummy (or phantom) filled with the reference material and the presence of air on both sides of the sample during the whole scan (Fig. 2F). For this reason a pipette tip was filled with FAA+PTA, sealed on the bottom end with paraffin wax, shortened at the top and sealed with parafilm (Fig. 2F). For one dummy scan to be used on sample scans, the following parameters have to remain constant: number of projection images, voltage, objective type, beam hardening coefficient, and source filter ((Candell, 2009); Fig. 2G). In practice, for all the scans in which we used CT scaling, projection images, voltage, objective type, beam hardening coefficient, and source filter were constant, whereas only voltage varied (Table S2). We thus carried out a dummy scan for each voltage value.

Calibration was performed as described in the manual (Candell, 2009). The density of FAA+PTA was measured seven times by using 4x a 100ml and 3x a 50ml measuring cylinder and an arithmetic mean of 93.7g/100ml was determined and used for calibration. Pollinia were reconstructed with CT scaling.

### Data processing

Two different procedures were developed to count pollen (summarised in Fig. 3A-D) and ovules (summarised in Fig. 3E-I). The pollen count procedure involves a high and a low resolution scan. In the low resolution scan, the whole pollinium is visible so that whole pollen volume is accessible. However in the low resolution scan, individual pollen grains are not resolved due to the strong aggregation of the grains. A high resolution scan (Fig. 3B) is thus used to estimate the number of pollen grains in a domain of the low resolution scan (Fig. 3C), which is then extrapolated to the whole pollinium (Fig. 3D). The estimation of the pollen grain number in the high resolution scan is performed *via* a series of thresholding and image processing steps followed by automatic object counting (summarised in Fig. 4, and detailed below).

**Fig. 3.**
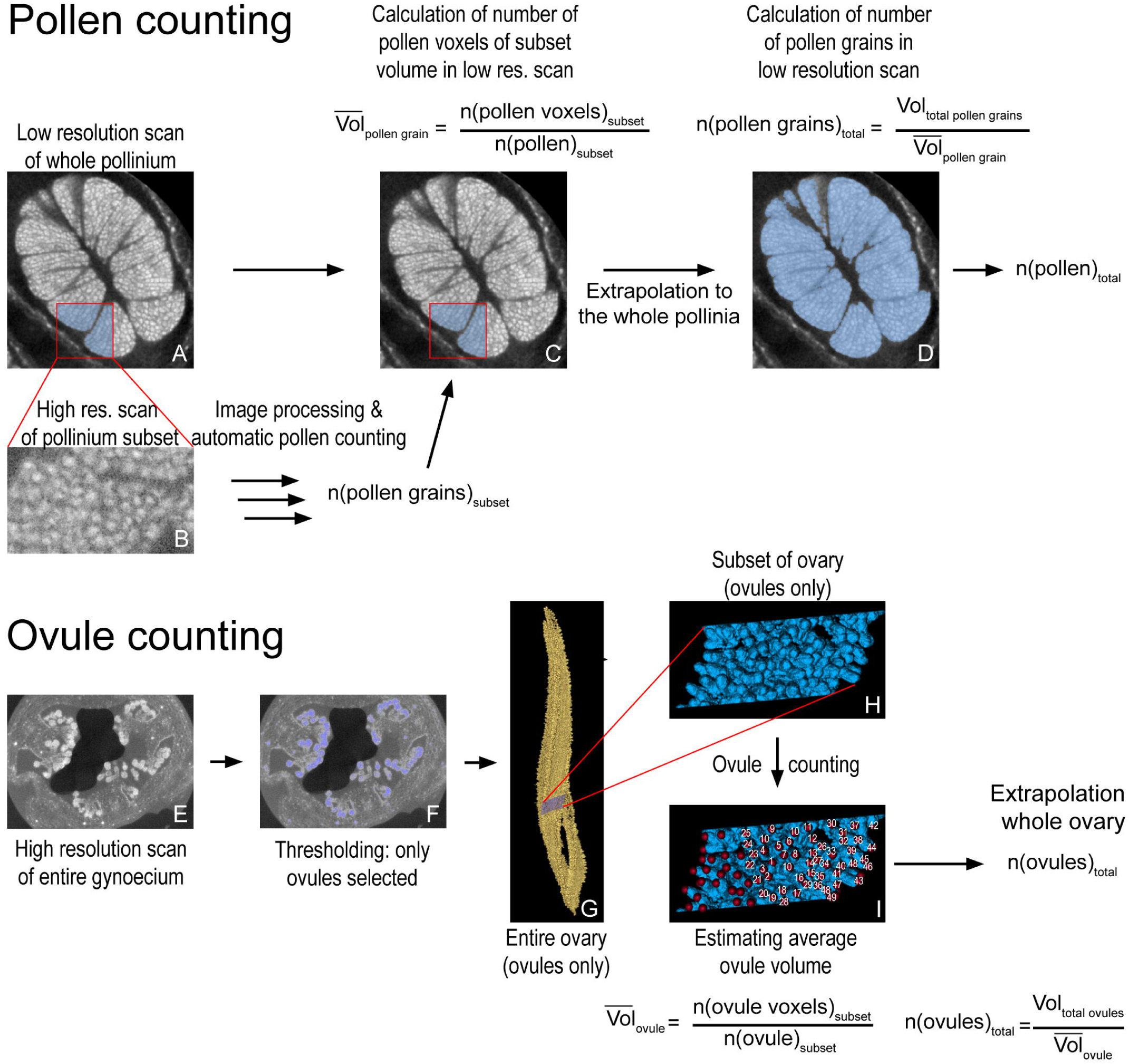
Workflows for object counting. A-D Workflow for counting object when individual objects cannot be resolved on a scan of the whole tissue, e.g. pollen in orchid pollinium. A, reconstructed section through pollinium with subset area highlighted in blue. B, reconstructed section of high resolution scan of the subset area (raw data). After image processing and automated object counting, the number of grains in the subset is calculated. C, the number of pollen grain in the subset is used to calculate the average volume of a pollen grain in the overview scan. D, the average volume of a pollen grain in the overview scan is used to calculate total pollen grain number. E-I Workflow for counting object when individual objects can be resolved on a scan of the whole tissue, e.g., ovules in orchid ovary. E, reconstructed section through ovary. F, thresholding of ovules in ovary (section). G, thresholding of ovules in ovary (3D model), with subset highlighted in blue. H, 3D model of subset. I, counting of ovules in subset (using landmark function in AMIRA which registers how many points have been set). This count allows to estimate the average volume of a single ovule. This value is used to obtain total ovule number.

The images of the high resolution scan were imported into the data analysis software AMIRA 5.4.1 (Build 006-Se11b; Konrad-Zuse Zentrum Berlin (ZIB) and Visage Imaging Inc.). The raw data (Fig. 4A, B) was first thresholded (Fig. 4C, D), i.e. all data below a threshold of greyscale value was removed from the dataset. The data was then filtered *via* a 3D median filter (kernel size 3x3x3 voxels; see Fig. 4E, F), and a Gaussian smoothing filter (kernel size 9x9x9 voxels; Fig. 4G, H). The images were then exported as 3D TIFF files to Fiji (Schindelin *et al.*, 2012), a distribution of ImageJ (Rasband, 1997-2016). Single pollen grains were then separated *via* “3D Iterative Thresholding” ((Ollion *et al.*, 2013); Fig. 4I, J), thresholded (Fig. 4K, L) and counted with the “3D Object Counter” (Bolte and Cordelieres, 2006). Subsequently, the pollen volume of the subset of the high resolution scan was measured in the low resolution scan (*via* the cropping function; Fig. 3C). Finally, we calculated the number of pollen grains per pollen volume for the low resolution scan, and extrapolated that value to the whole flower (Fig. 3D), in order to obtain the total pollen grain number.

**Fig. 4.**
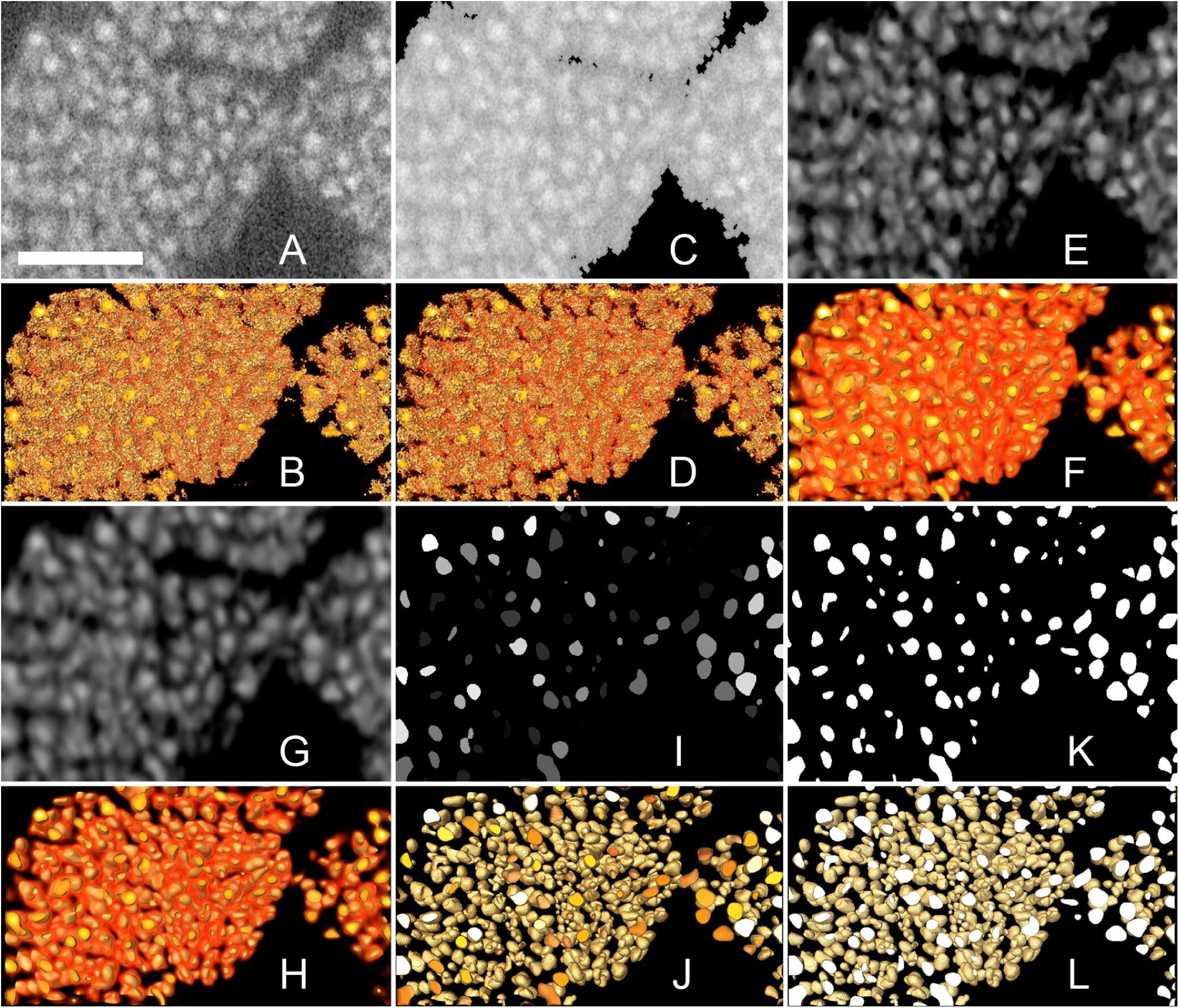
Image processing pipeline. Section through high resolution scan dataset of pollinium subset and associated 3D models illustrating the sequences of steps during image processing. A, reconstructed section of raw data. B, 3D model of the raw data. C, data after greyscale thresholding (removal of voxels darker than a specific value). D, 3D model of the data after greyscale thresholding. E, data after 3D median noise reduction filter. F, 3D model of data after 3D median noise filter. G, data after 3D Gaussian smoothing filter. H, 3D model of data after 3D Gaussian smoothing filter. I, data after iterative thresholding. J, 3D model of data after iterative thresholding. K, data after greyscale thresholding. L, 3D model of the data after greyscale thresholding. The number of objects in the scan data can now be automatically counted with the 3D Object Counter function of Fiji. Scale bar 50µm.

The ovule count procedure only involved one scan because the ovules can be distinguished on scans of the whole gynoecium. The ovules were segmented away from the rest of the ovary (Fig. 3E-G). On a subset of the ovules (Fig. 3H), the ovules were counted manually by using the landmark function (Fig. 3I). Average ovule volume was thereby calculated. The total ovule volume was then divided by the average ovule volume in order to obtain total ovule number per flower.

### Statistical tests

We investigated differences in the number of ovules, the number of pollen grains, and the value of the pollen to ovule ratio (P:O), between rewarding and deceptive species, among species, and within inflorescences. We therefore tested the effects of the factors “interaction” (deceptive and rewarding), “species” and “flower position” (bottom, middle, and top) on the number of ovules, on the number of pollen grains, and on the P:O per flower, respectively. Analyses were performed with the software R (R_Core_Team, 2014). Because our data is nonparametric (see Supplementary data S1), we used a non-parametric analysis of variance (npANOVA), with the function Adonis (vegan) (Oksanen *et al.*, 2013). We first generated a distance matrix with the function dist(stats) using the Euclidean distance index, and then performed the npANOVA using 9999 permutations (Anderson, 2001). Post hoc tests were performed with the same function, with a Bonferroni correction for multiple comparisons. Finally, we used the same test to look for an effect of the interaction between the factors “species” and “flower position”. If the effect between the factor “species” and “flower position” is significant, it means that the number of pollen or ovules does not vary in the same direction within the inflorescence for each species. If this effect is not significant, it means that the number of ovules or pollen grains varies in the same direction within the inflorescences for all species. The variance of the P:O per species was compared between rewarding and deceptive species with a non-parametric Mann-Whitney-Wilcoxon (MWW) test using the function wilcox.test(stats).

### Phylogenetic analyses

We used a pruned version of the time-calibrated phylogeny of Inda et al. (Inda et al., 2012) for phylogenetic comparative analyses (Fig. 5E). We did not include *Dactylorhiza majalis* in our phylogeny, because of its allotetraploid origin (Hedren et al., 2001). First, we calculated the amount of phylogenetic signal in individual traits using the maximum-likelihood value of Pagel’s λ (Pagel, 1999). We estimated Pagel’s λ using the function fitContinuous based on likelihood optimisation (ML) for four continuous traits and fitDiscrete based on symmetric model (SYM) for one discrete trait, in the GEIGER package (Harmon et al., 2008). λ = 0 indicates that there is no phylogenetic signal for the trait, which means that the trait has evolved independently of phylogeny, i.e. close taxa are not more similar on average than distant taxa. λ = 1 indicates that there is a strong phylogenetic signal, which means that the trait has evolved according to the Brownian motion model of evolution (Pagel, 1999).

**Fig. 5.**
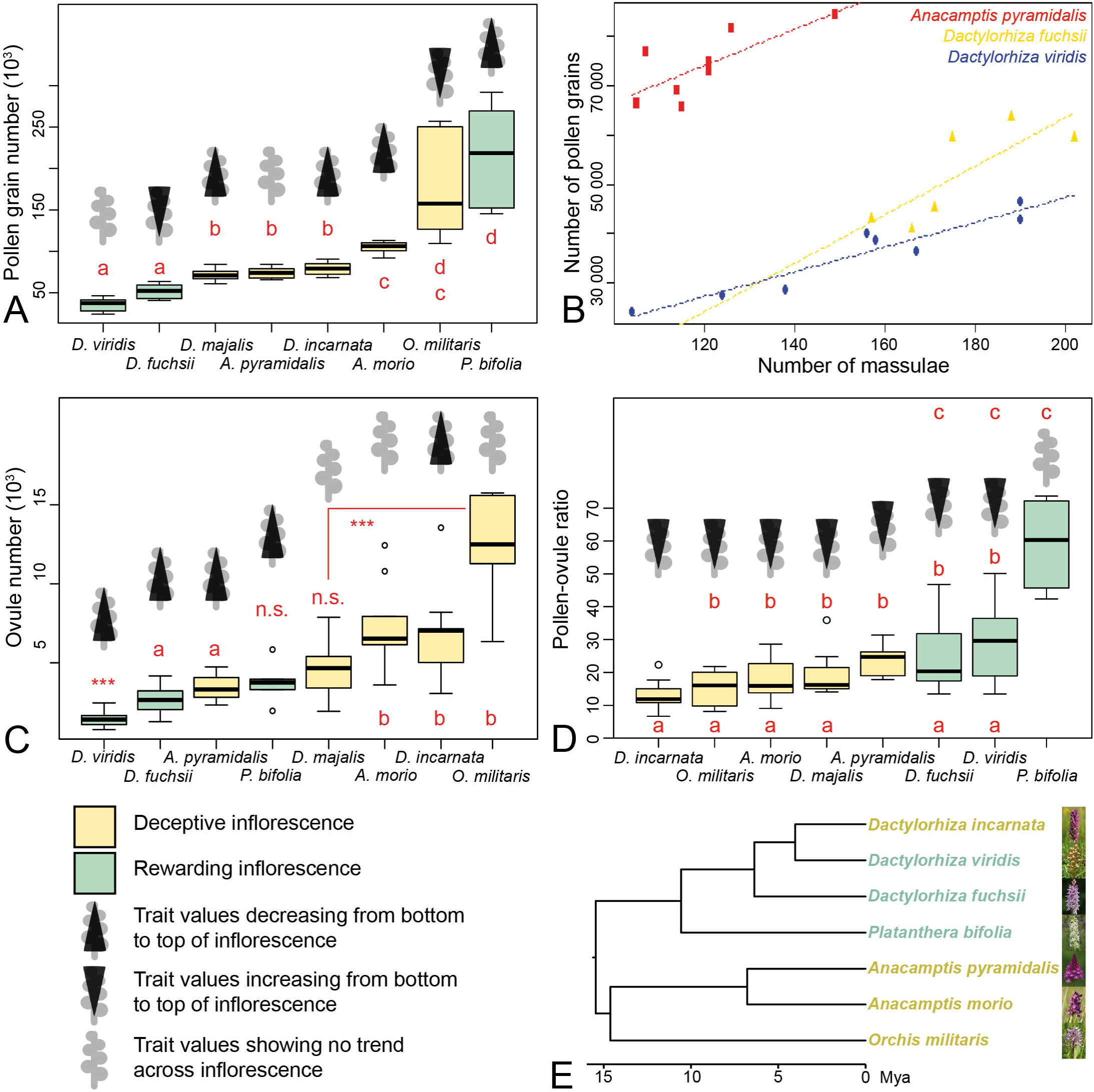
Pollen, massulae, ovules, pollen:ovule ratio, and phylogeny. A, pollen grain number per flower for the eight orchid species studied. B, massulae number per flower and corresponding pollen grain numbers for a subset of three species. C, ovule number per flower for the eight orchid species studied. D, P:O per flower for the eight orchid species studied. E, time calibrated phylogeny of the eight orchid species studied. A-D, data displayed in boxplot format. Letters indicate species that group together according to the npANOVA posthoc tests. *** = group that differs from all the others, or significant difference between two groups; n.s. = group that differs from none of the others

We also applied phylogenetic generalised least squares (PGLS; (Martins and Hansen, 1997)) to understand the nature of the evolutionary association between biological traits, as implemented in the R package caper (Orme, 2013). We identified one predictor variable (presence/abscence of floral reward) that could affect the response variables (pollen grain number, ovule number, P:O, and variance thereof) and ran a PGLS including all the variables. First, a variance–covariance matrix was calculated based on the phylogenetic relationships of the species. In the PGLS, λ applies to the residual errors from the regression model, not to the strength of the signal in the response variable, nor to that of the predictor variables. λ = 0 indicates a non-phylogenetic covariance matrix, whereas λ = 1 refers to the expected phylogenetic covariance matrix under a Brownian motion model of evolution (Garamszegi, 2014).

## Results

### Development of a new method

We present a set of two approaches to count numerous high contrast objects in plant tissues *via* X-Ray CT. One method is presented for cases where it is possible to resolve all the objects to be counted on a scan of the whole tissue. The other method is applicable when this is not the case. Ovules can be resolved on scans of the whole ovary; therefore, an estimation of the average volume of an ovule and an ovule count can be carried out on the same data (Fig. 3E-I). At least in the special of case of Orchidaceae, where pollen grains are relatively small and aggregated in compact pollinia, individual pollen grains cannot be resolved on the same scan data as overview scans of the entire pollinium. Two scans are therefore necessary (see Fig. 3A-D). A high resolution scan of a subset of the tissue has to be performed in order to estimate the number of pollen grains inside this subset (Fig. 3B, C); this number will then be used to estimate total object number on the overview scan (Fig. 3D). On the high resolution scan, image processing and automatic counting methods are presented that allow the straightforward processing of scan data (Fig. 4).

### Variation of number of pollen grains and massulae per flower

Pollen grain number per flower significantly differed among species (npANOVA: F = 34.32; R^2^ = 0.81, p = 0.001; Table 1) and ranged from 35 588 ± 2 823 in *Dactylorhiza viridis* to 215 978 ± 25 687 in *P. bifolia*. There was no significant difference in pollen grain number per flower between rewarding and deceptive species (npANOVA: F = 0.02; R^2^ = 0.00, p = 0.918). The number of pollen grains per flower did not significantly differ within inflorescences (npANOVA: F = 0.7; R^2^ = 0.01, p = 0.520). This lack of significance is likely due to our small sample size. In fact, when looking at the data, pollen grain number tended to increase from bottom to top in *Orchis militaris* and *D. fuchsii*, and to decrease from bottom to top in *P. bifolia*, *D. majalis*, *D. incarnata*, and *A. morio*. There was no trend of variation in *D. viridis* and *A. pyramidalis* (Fig. 5A).

**Table 1.** **Mean pollen and ovule numbers, pollen to ovule ratio and variance thereof in the eight species studied and comparison with previously published values** Mean low. = mean number of ovules on bottom flowers of inflorescences. Se = standard error. N = sample number. Var = variance. lit. = literature. a (Neiland and Wilcock, 1995); b* (Salisbury, 1942);c* (Jersáková and Kindlmann, 1998); d* (Nazarov, 1995); e* (Na zarov, 1998); f*(Darwin, 1862); g (Nazarov and Gerlach, 1997); h (Daumann, 1941); i* (Claessens and Kleynen, 2011); j (Dafni and Woodell, 1986); k (Inda *et al.*, 2012) and references therein; l (Hansen and Olesen, 1999); m* (Sonkoly *et al.*, 2016). * Ovules counted as seeds.

| Species | Pollen grain number |  |  |  | Ovule number |  |  |  |  | Pollen : Ovule ratio |  |  |  | Deceptive<br>Rewarding |
| --- | --- | --- | --- | --- | --- | --- | --- | --- | --- | --- | --- | --- | --- | --- |
|  | mean | Se | N | from lit. | mean | Se | mean low. | N | from literature | mean | Se | Var | from lit. |  |
| *Anacamptis morio* | 104817 | 3207 | 6 | - | 7347 | 952 | 9827 | 9 | >4000 b, 4770 ± 1856 c<br>5052 d, 4978 ± 521 m | 18 | 2.9 | 49 | 13 a | D k |
| *Anacamptis pyramidalis* | 74013 | 2407 | 8 | - | 3435 | 232 | 4350 | 11 | 1935 b, 3036 d<br>2262 ± 205 m | 24 | 1.7 | 22 | - | D h |
| *Dactylorhiza fuchsii* | 51917 | 4075 | 6 | - | 2659 | 327 | 3413 | 8 | 6200 f, 3294 ± 774 b<br>5205 ± 914 m | 25 | 5.0 | 152 | 21 a | R j |
| *Dactylorhiza incarnata* | 79113 | 2761 | 8 | 194748 g | 6715 | 846 | 9030 | 11 | 7270 d, e 7756 g<br>7076 ± 881 m | 13 | 1.7 | 23 | 12.6 (25.2) g | D k |
| *Dactylorhiza majalis* | 72050 | 2422 | 10 | - | 4528 | 455 | 5447 | 12 | 9639 ± 421 m | 19 | 2.2 | 46 | - | D l |
| *Dactylorhiza viridis* | 35588 | 2823 | 8 | - | 1413 | 134 | 1772 | 12 | 1339 ± 693 b<br>1453 ± 136 m | 29 | 4.3 | 147 | - | R i |
| *Orchis militaris* | 176417 | 26062 | 6 | - | 12616 | 1296 | 14183 | 7 | 10948 ± 3274 m | 15 | 2.2 | 30 | - | D k |
| *Platanthera bifolia* | 215978 | 25687 | 6 | 217396 g | 3779 | 514 | 4930 | 6 | 3666 d, e 4004 g<br>6146 ± 325 m | 59 | 5.4 | 175 | 27.1 (54.3) g | R k |

The number of massulae (sub-aggregates within the pollinium) per flower was determined for *Anacamptis pyramidalis*, *D. fuchsii*, and *D. viridis* (see table S3). The number of massulae per flower ranged in *A. pyramidalis* from 105-149, in *D. fuchsii from* 157-202, and in *D. viridis* from 104 to 190. A positive correlation between number of massulae per flower and the total number of pollen grains could was present (Fig. 5B). In two species this correlation was significant: in *A. pyramidalis* (F_1,6_=7.77, p=0.03168, R^2^=0.4916), and in *D. viridis* (F_1,6_=56.57, p=0.0.000286, R^2^=0.8881), whereas in *D. fuchsii* only a trend could be detected (F_1,4_=6.942, p=0.0579, R^2^=0.543).

### Variation of ovule number per flower

Ovule number per flower significantly differed among species (npANOVA: F = 29.3; R^2^ = 0.45, p = 0.001; Fig.5C) and ranged from 1,413 ± 134 in *D. viridis* to 12 616 ± 1 296 in *O. militaris* (Table 1). Deceptive species produced in average significantly more ovules per flower (6 408 ± 521) than rewarding species (2 342 ± 248; npANOVA: F = 112.5; R^2^ = 0.29, p = 0.001). Finally, ovule number per flower tended to decrease from bottom to top of the inflorescences for all species (“flower position”: npANOVA: F=19.5; R^2^ = 0.10, p = 0.001; effect of the interaction “species” × “flower position”: npANOVA: F=1.3; R^2^ = 0.04, p = 0.256). This trend was not clear for *A. morio*, *D. majalis* and *O. militaris* (Fig. 5C).

### Pollen/ovule ratio

The P:O per flower significantly differed among species (npANOVA: F = 16.28; R^2^ = 0.36, p = 0.001; Fig. 5D) and ranged from 13.06 ± 1.70 in *D. incarnata* to 59.08 ± 5.41 in *P. bifolia* (Table 1). The P:O in rewarding species (36.91 ± 4.27) was twice higher than in deceptive species (17.95 ± 1.08; npANOVA: F = 112.5; R^2^ = 0.29, p = 0.001), and in average more than four times more variable (MWW-test: n_1_ = 3; n_2_ = 5; W = 0; p = 0.035, Table 1). Finally, the P:O tended to increase from bottom to top of the inflorescences for all species (“flower position”: npANOVA: F = 17.80; R^2^ = 0.12, p = 0.001; effect of the interaction “species” × “flower position”: npANOVA: F = 0.91; R^2^ = 0.05, p = 0.545). This trend was not clear for *P. bifolia* (Fig. 5D).

### Phylogenetic analyses

Individually, the traits used in the analyses exhibit a λ nearly always zero, indicating no phylogenetic signal except for the “mean pollen number” (λ=1; see Table S4). In contrast to the phylogenetic signal estimates for the individual variables, the estimated maximum likelihood values of λ for two of the regression models (“mean ovule number” ˜ “pollination strategy” (deceptive or rewarding) and “P:O variance” ˜ “pollination strategy”) were zero, indicating no phylogenetic signal in the residual errors of the models, and hence results that are equivalent to conventional ordinary least squares analyses. However, two other regression models (“mean pollen grain number” ˜ “pollination strategy” and “P:O” ˜ “pollination strategy”) were under a Brownian motion model of evolution with an estimated maximum likelihood value of λ = 1. The “pollination strategy” was only strongly associated with the “P:O variance” (table S5).

## Discussion

### Development of a new method

Traditionally, pollen and ovule counts have relied on destructive sampling and sub-sampling. For example, in orchids pollen counts have relied on pollen counts of single massulae (sub-aggregates within the pollinium) which are then extrapolated to the whole pollinia (Nazarov and Gerlach, 1997).

This method is flawed for two reasons: (1) within a pollinium, massulae have widely different numbers of pollen grains; often there are a few large massulae and many smaller ones. These differences in size have been shown to be stronger in deceptive orchids than in rewarding ones (Nazarov and Gerlach, 1997) This unequal distribution is probably the reason why our counts of pollen grains in the rewarding *Platanthera bifolia*, which has many massulae of similar size, is close to the pollen counts published by (Nazarov and Gerlach, 1997), whereas in the deceptive *Dactylorhiza incarnata*, the pollinia of which are composed of massulae of widely different sizes, our values are very different from (Nazarov and Gerlach, 1997). Nazarov and Gerlach (1997) also possibly over-estimated the numbers of pollen grains in *D. incarnata* because by counting the grains mostly in the few, large massulae, they probably overestimated the contribution of the many small massulae. (2) extrapolating values from massulae assumes that, within the same species, massulae numbers and pollen grain numbers are always tightly correlated, which we show not to be correct (see Fig. 5B). Counting methods relying on the whole volume of pollen are therefore more precise than methods relying on counts of massulae. Our methods also do not require de-aggregation of the pollen grains. Pollen aggregation evolved at least 39 times in angiosperms, including in some of their most species-rich lineages (e.g. the legumes and the orchids), probably because it promotes male fitness in response to infrequent pollinator visits (Harder and Johnson, 2008). The counting method we present allows to bypass the requirement of de-aggregation of the pollen grains, and can provide accurate counts for any type of pollen.

For the species we studied, the seed counts from the literature were mostly obtained from flowers that were naturally pollinated (see table 1 and citations therein). We therefore assume that, in species with deceptive flowers, only 10% of all fruits came from flowers from the top third of the inflorescence. In species with rewarding flowers, we assume that the seeds were counted from all portions of the inflorescences equally. With these assumption our counts are on average 4,5% higher than the values from literature (see table 1 and citations therein), although variation is very large and makes comparisons difficult. Our higher counts are probably due to our method counting all ovules, not only those that were fertilized. We could not test the latter possibly due to the specificities of orchid seed development (see Supplementary data S2). In the gynoecia of flowers at the top of inflorescences, extensive gaps in the ovule distribution were often noticed (Fig. S1 A-E). These gaps could affect the chemotactic signalling from ovules to guide pollen tubes (Okuda *et al.*, 2009; Takeuchi and Higashiyama, 2016), which could lead to some ovules not being fertilized, even if the pollen loads would be sufficient to do so.

And finally, unlike all the previously existing methods that require the destruction of the sample, our processing methods allow to measure both pollen and ovule numbers from the same flower with minimal destruction (perianth removal). The column and ovary are left intact after counting, and could be used for other analyses: shape analysis, histology, etc. The methods we present are flexible and could be used to count any high contrast objects inside plant tissues, such as crystals (druses or raphides) or stone cells (sclereids).

### Reproductive investment and presence/absence of floral reward

There is no significant difference in pollen grain number per flower between rewarding and deceptive species, probably because the two rewarding species *D. viridis* and *D. fuchsii* produced significantly less pollen than all the other species, whereas the third rewarding species, *P. bifolia*, was one of the species producing the most pollen grains per flower (Fig. 5A). There are also no significant trends of increase or decrease across inflorescences. Moreover, pollen grain number appears to follow a Brownian Motion model of evolution. Taken together these data suggest that pollen grain number is not under strong selection.

Ovule numbers strongly differ between species with rewarding and deceptive flowers. Deceptive flowers contain more ovules than rewarding flowers ((Sonkoly *et al.*, 2016); this study), which is possibly an adaptation that enables deceptive inflorescences to have the same seed set as rewarding inflorescences, despite lower fertilization rates (Sonkoly *et al.*, 2016). Ovule numbers do not show any phylogenetic signal, possibly due to high homoplasy in this character state, i.e. rapid evolution in adaptation to different reproductive ecologies. In both rewarding and deceptive species, ovule numbers decreased from bottom to the top of the inflorescences.

Due to lack of difference in pollen grain numbers and strong difference in ovule numbers across pollination strategy and positions in inflorescence, differences in P:Os are driven by ovule numbers. Increase in P:Os from bottom to top of inflorescences have been observed in orchids (Kopylov-Gus’ kov Yu *et al.*, 2006; Nazarov and Gerlach, 1997; Salisbury, 1942), and other taxa, see e.g. (Thomson, 1989). Significant differences in variance of P:O between deceptive and rewarding flower and the well-supported strong phylogenetic correlation of these two characters (P:O variance and pollination strategy) highlight the differences in constraints acting on the reproductive investment in deceptive *versus* rewarding species. Given that deceptive flowers contain significantly more ovules than rewarding flowers ((Sonkoly *et al.*, 2016); this study), and given that usually only the lower flowers on the inflorescences of deceptive flowers are fertilized (Jersáková and Kindlmann, 1998; Nilsson, 1980; Vogel, 1993), it seemed likely that deceptive inflorescences are under strong constraint to stringently decrease ovule numbers from bottom to top of the inflorescence in order to efficiently allocate resources. In rewarding flowers there are much fewer ovules, and fertilization occurs across the whole inflorescence. It seems thus likely that constraints acting on reproductive allocation in rewarding inflorescences is much weaker, which could explain the much larger variance in P:Os found in rewarding inflorescences.

Ultimately, the new tools we present will allow much broader comparative studies of the diverse groups of angiosperms in which pollination by deceit takes place, and will allow us to better quantify plant reproductive investment.

## Supplementary data

### Supplementary data S1

Choice of ANOVA test

### Supplementary data S2

Loss of contrast of developing seeds in selected European orchid species and comparison with *Arabidopsis thaliana*

**Table S1** Collected species, locality, collection date, breeding strategy, pollinators, and permits.

**Table S2** Computed Tomography scanning parameters. For all scans, camera binning = 1; top = flower from top portion of the inflorescence; mid = flowers from middle portion of the inflorescence; bot = flowers from the bottom part of the inflorescence; pol = pollinia; gyn = gynoecium.

**Table S3** Counts of pollen grains, massulae, and ovules. For pollen number, N/A was reported when the pollinium lost massulae during preparation (total pollen grain number was not available). Massulae numbers were counted for *Anacamptis pyramidalis*, *Dactylorhiza fuchsia*, and D. viridis (other species denoted with N/A)

**Table S4.** Phylogenetic signal estimates maximum likelihood values of Pagel’s lambda for continuous traits and symmetric model for a discrete trait.

| Continuous Traits | Maximum likelihood values of Pagel's lambda |
| --- | --- |
| Mean Pollen Grain number | 1 |
| Mean ovule number | 0.29 |
| Pollen : Ovule ratio | 1.082E-211 |
| Variance Pollen : Ovule ratio | 0.08459718 |
| Discrete Trait | Symmetric model values of Pagel's lambda |
| Pollination strategy (Deceptive / Rewarding) | 3.56E-115 |

**Table S5.** Phylogenetic Generalized Least Squares Analysis for four models - coefficient for predictors.

| Model components | Predictor of pollination strategy | Pagel's lambda | Adjusted R<sup>2</sup> (%) | Coefficient for predictor | p value |
| --- | --- | --- | --- | --- | --- |
| Mean pollen grain number<br>~ pollination strategy | Mean pollen grain number | 1 | 1.86 | 0.66 | 0.3395 |
| Mean ovule number<br>~ pollination strategy | Mean ovule number | 0 | 4.82 | 1.49 | 0.04823 |
| Pollen : ovule ratio<br>~ pollination strategy | Pollen : ovule ratio | 1 | 67.79 | -1.17 | 0.01413 |
| Variance pollen : ovule ratio<br>~ pollination strategy | Variance pollen : ovule ratio | 0 | 90.35 | -2.42 | 0.00064 |

**Movie S1** 3D models of pollinium subset high resolution scan data after each steps of image processing, from raw reconstructed data to machine countable objects.

## Acknowledgements

The authors would like to thank Manuel Pimentel and Luis Ángel for sharing their phylogenetic tree file. The authors would like to thank Manfred Fischer’s whose encyclopedic knowledge of the Austrian flora made this project possible. Luise SchrattEhrendorfer- and Harald Niklfeld for their help to plan and carry out the field work.

---

**Fig. S1.**
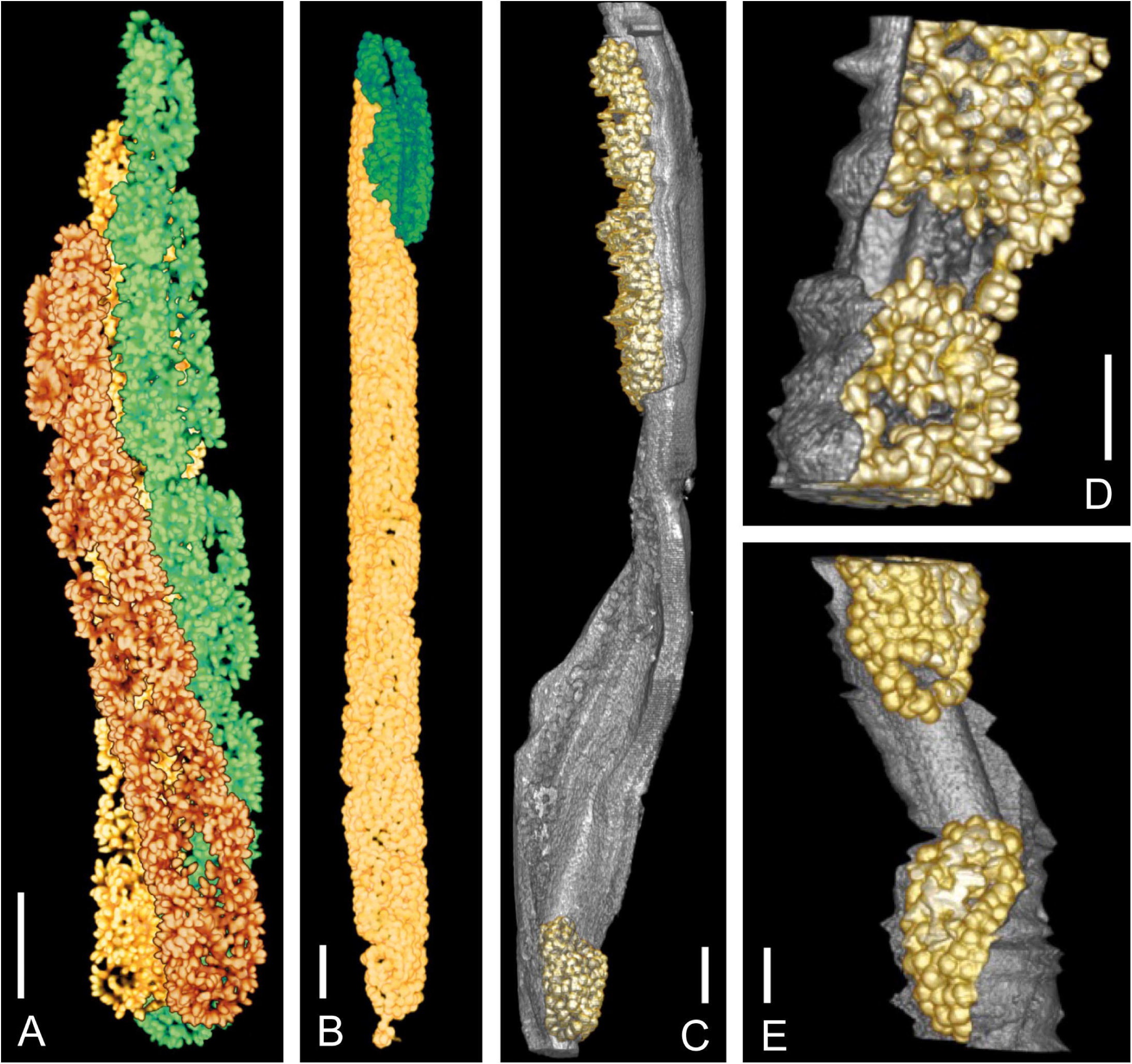
Ovule distribution inhomogeneity. Ovule distribution differences between flowers from the bottom of the inflorescence and flowers from the top of the inflorescence. Ovules from different placentae are in different colours; placentae are in grey; distal end of gynoecia towards the top of the page. A, ovules from gynoecium of bottom flower of *Dactylorhiza incarnata*. B, top flower of *D. majalis*, one placenta is completely missing, and one is present only for less than 25% of the length of the ovary (in green). C, flower from top of inflorescence of *D. majalis*; more than 50% of the placenta length is devoid of ovules. D, flower from top of inflorescence in *Anacamptis pyramidalis*, small gap of ca. 5% of placenta length. E, flower from top of inflorescence in *D. incarnata*, small gap of ca. 14% of placenta length. Sc

